# The global anaerobic metabolism regulator *fnr* is necessary for the degradation of food dyes and drugs by *Escherichia coli*

**DOI:** 10.1101/2022.10.25.513779

**Authors:** Lindsey M. Pieper, Peter Spanogiannopoulos, Regan F. Volk, Carson J. Miller, Aaron T. Wright, Peter J. Turnbaugh

## Abstract

The microbiome is an underappreciated contributor to intestinal drug metabolism with broad implications for drug efficacy and toxicity. While considerable progress has been made towards identifying the gut bacterial genes and enzymes involved, the role of environmental factors in shaping their activity remains poorly understood. Here, we focus on the gut bacterial reduction of azo bonds (R-N=N-R’), found in diverse chemicals in both food and drugs. Surprisingly, the canonical *azoR* gene in *Escherichia coli* was dispensable for azo bond reduction. Instead, azo reductase activity was controlled by the fumarate and nitrate reduction (*fnr*) regulator, consistent with a requirement for the anoxic conditions found within the gastrointestinal tract. Paired transcriptomic and proteomic analysis of the *fnr* regulon revealed that in addition to altering the expression of multiple reductases, FNR is necessary for the metabolism of L-Cysteine to hydrogen sulfide, enabling the degradation of azo bonds. Taken together, these results show how gut bacteria sense and respond to their intestinal environment to enable the metabolism of chemical motifs found in both dietary and pharmaceutical compounds.

**IMPORTANCE:** This work has broad relevance due to the ubiquity of dyes containing azo bonds in food and drugs. We report that azo dyes can be degraded by human gut bacteria through both enzymatic and non-enzymatic mechanisms, even from a single gut bacterial species. Furthermore, we revealed that environmental factors, oxygen and cysteine, control the ability of *E. coli* to degrade azo dyes due to their impacts on bacterial transcription and metabolism. These results open up new opportunities to manipulate the azoreductase activity of the gut microbiome through the manipulation of host diet, suggest that azoreductase potential may be altered in patients suffering from gastrointestinal disease, and highlight the importance of studying bacterial enzymes for drug metabolism in their natural cellular and ecological context.

## INTRODUCTION

While it has long been appreciated that the gut microbiota, the trillions of microorganisms found in the gastrointestinal (GI) tract, and its aggregate genomes (the gut microbiome) contribute to the digestion and metabolism of dietary macronutrients, the broader role of the microbiome in the metabolism of xenobiotics (diet-derived and pharmaceutical small molecules) is less well understood. Recently, work in cell culture, mice, and humans have emphasized that both excipients (food and drug additives) and pharmaceuticals are extensively metabolized by the gut microbiota (1–6). The presence of an azo bond (R-N=N-R’) is shared between both pharmaceuticals and excipients (7, 8), which is notable given the ability of diverse human gut bacteria to reduce azo bonds (9). While azo reductase activity may be a core metabolic function of all human gut microbiotas (10–14), the enzymatic and non-enzymatic mechanisms responsible for this activity and their sensitivity to environmental factors remain poorly understood.

Mechanistic insights into this process are essential given the broad impact of azo reduction for antibiotics (15) and anti-inflammatory drugs (7, 16). Furthermore, the consumption of food, drug, and cosmetic (FD&C) dyes is increasing (17), providing additional substrates for gut bacterial metabolism. We recently discovered that FD&C dyes are potent inhibitors of the influx transporter OATP2B1, interfering with drug absorption in gnotobiotic mice (18). This effect was rescued by human gut bacterial metabolism due to an inability of the downstream microbial metabolites to inhibit OATP2B1 (18). Additional work in rodent models has implicated FD&C dyes in carcinogenesis (19–21) and inflammatory bowel disease (22). Thus, the ability to predict or control azo reductase potential within the human gut microbiome could have broad implications for host health and disease.

The canonical enzyme implicated in this metabolic activity is azo reductase (AzoR) (9, 11, 14, 23). Work on the model human gut bacterium *Escherichia coli* has demonstrated that the purified AzoR protein is sufficient to reduce azo bonds (24). For example, *Pseudomonas aeruginosa* (25–28)and *Enterococcus faecalis* (29–31)encode multiple azo reductases. Alternative mechanisms have been proposed for azo reductase activity, including the electron transport chain of *Shewanella oneidensis*,nicotinamide adenine dinucleotide (NADH), and hydrogen sulfide (H_2_S) (32–34). While these studies provide valuable mechanistic insight, a major limitation is their focus on *in vitro* biochemistry, neglecting to address the cellular and genetic mechanisms that impact azo reductase activity or the potential confounding effects of the complex physiological and microbiological interactions within the GI tract.

We sought to address this knowledge gap through the mechanistic dissection of a representative member of the human gut microbiota. Surprisingly, we found that the *azoR* gene is dispensable for the azo reductase activity of *E. coli*. In our search for alternative azoreductase enzymes, we discovered that *E. coli* azo reduction is regulated by the oxygen-sensing dual-transcriptional regulator, FNR. In turn, we discovered that FNR regulates L-Cysteine metabolism, which produces H_2_S. Surprisingly, *E. coli*-produced H_2_S is sufficient to reduce azo dyes and drugs. Taken together, we demonstrate that environmental oxygen regulates the metabolism of L-Cysteine to H_2_S which degrades commonly consumed azo-bonded compounds. Our results highlight the importance of studying and developing an understanding of the environmental context and regulation of key microbial metabolisms beyond annotated enzyme function.

## RESULTS

### *E. coli* AzoR is dispensable for azoreductase function

Prior work from our lab (18, 35) and others (36) has identified widespread homologous genes to *E. coli* azo reductase (AzoR) in the genomes of human gut bacteria capable of metabolizing diverse compounds containing azo bonds. We had originally sought to establish a tractable system to study these heterologously expressed genes by first abolishing this activity in *E. coli*. Much to our surprise, the *E. coli* Keio collection azoreductase knockout strain (Δ*azoR*) had activity indistinguishable from wild-type (*wt*) during growth in both solid (**Figure 1A**) and liquid (**Figure 1B**) media containing a representative azo bond-containing food coloring (FD&C Red No. 40). We validated this finding by constructing and testing a clean deletion of *azoR* (**Figure 1C**), which was confirmed by gel electrophoresis (**Figure S1A**) and Sanger sequencing (**Figure S1B**).

**FIG 1.**
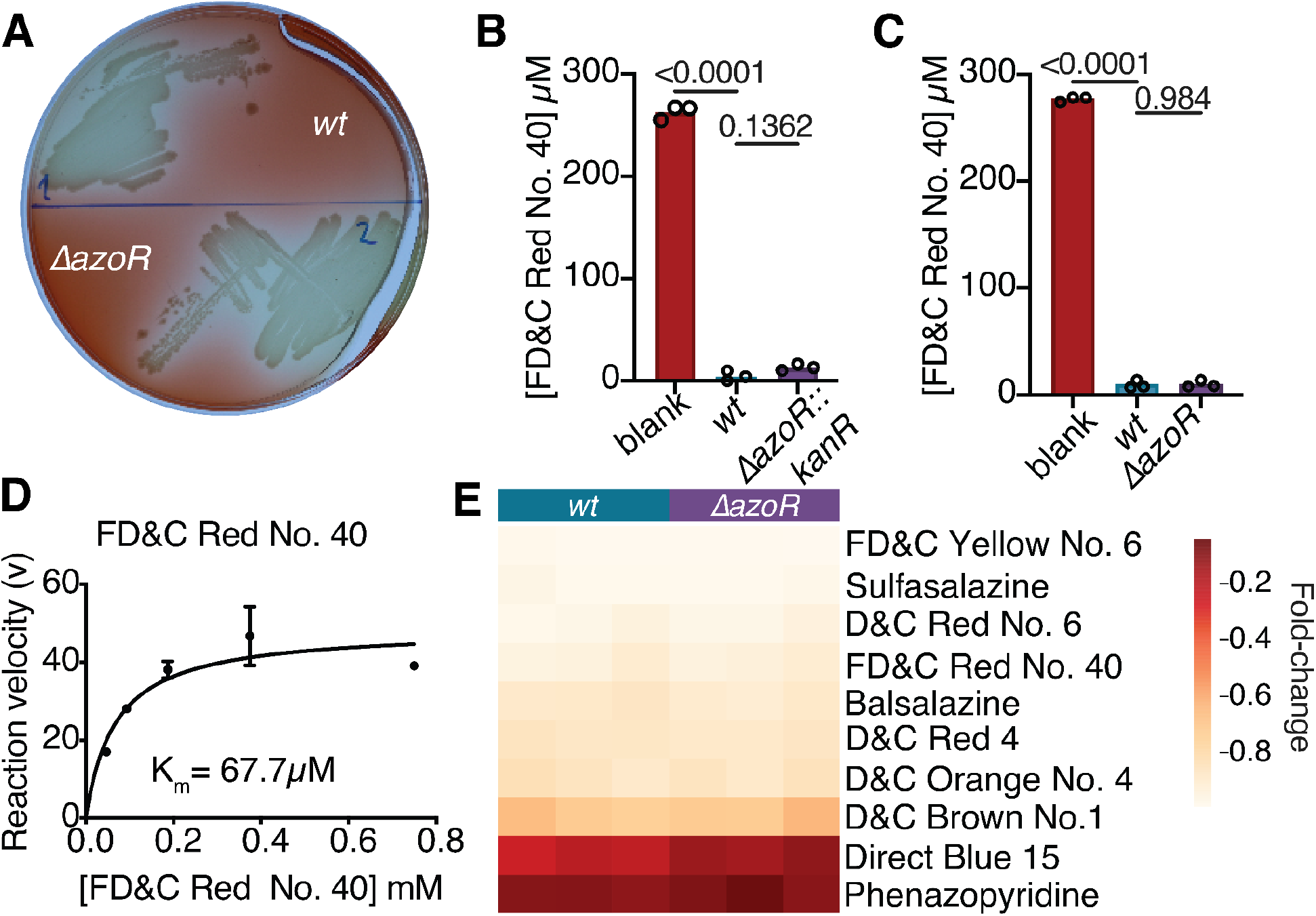
*E. coli* AzoR is sufficient but dispensable for azoreductase activity. *E. coli azoR* is dispensable for azo reductase activity during anaerobic growth on (A) solid and (B) liquid media. (C) knockout phenotype holds with a clean *azoR* deletion. (A-C) 25 μM azo dye/drug. (A) 72 hours growth on LB agar with 4.13 mM L-Cysteine. (B,C) 24 hours growth in LB with 4.13 mM L-Cysteine. Supernatant was removed from samples and analyzed for residual dye concentration spectrophotometrically. Data are normalized to uninoculated controls. *p*-values, one-way ANOVA. (D) Michaelis-Menten curve for purified AzoR enzyme with FD&C Red No. 40. Michaelis-Menten curve fitted using PRISM. E) Dye depletion in liquid media by *wt* or *ΔazoR* across a panel of azo drugs and dyes. Grown with 250 μM azo dye/drug, for 24 hours in LB with 4.13 mM L-Cysteine. Fold-changes are relative to average blank concentration (n=3 biological replicates/strain).

Given these unexpected results, we wondered if AzoR might be inactive in the tested *E. coli* strains, potentially due to mutations that could have occured during the creation of our lab’s culture collection (37). Sanger sequencing of the *azoR* gene verified that the coding sequence was 100% identical to the deposited *E. coli* BW25113 genome. To further validate the activity of this enzyme, we purified the Histidine-tagged AzoR protein from *E. coli* AG1 with the pCA24N plasmid for overexpression (ASKA collection) (38). Cells were grown to mid-exponential growth phase, *azoR* expression was induced with Isopropyl β-d-1-thiogalactopyranoside (IPTG), then cells were harvested, and protein was purified using a nickel-charged affinity column. The fraction location of the protein was determined using an SDS-PAGE gel and the size of the protein was consistent with AzoR (**Figure S2A**). As expected, the purified protein was sufficient to clear a panel of azo bond containing dyes and drugs (**Figure S2B)**. Additionally, we were able to create a Michaelis-Menten curve describing the enzymatic activity against FD&C Red No. 40, where we calculated a K_M_ of 67.7μM (**Figure 1D**). Finally, we tested our whole cell assay using a panel of 10 food additives and pharmaceutical compounds containing azo bonds, demonstrating that *azoR* is unnecessary for the depletion of multiple dyes (**Figure 1E**). These results confirm that the canonical AzoR enzyme in *E. coli* (24)is sufficient for azo reductase activity; however, *azoR* is not required for azo bond reduction by *E. coli* cells, emphasizing that biochemical activity of purified proteins is not necessarily predictive of metabolism in whole cells and motivating a renewed search for the mechanism(s) responsible.

### FNR is necessary for the *in vitro* azoreductase activity of *E. coli*

To identify genes necessary for azo dye depletion, we screened 115 candidate gene deletions from the Keio collection using a high-throughput liquid media assay for the depletion of the widely used food coloring FD&C Red No. 40. Target genes were chosen based on previously published literature. First, work with a transposon mutagenesis library of *Shewanella oneidensis* MR-1 demonstrated that the electron transport chain is involved in azo dye depletion (39). Homologs of these genes were identified in *E. coli* and tested (see *Methods*). Additionally, Zhang *et al*. tested gene expression in *E. coli* that were bound to a quinone and exposed to azo dyes (40). Genes that were upregulated when exposed to azo compound, annotated as involved in electron transport, or annotated as reductases were also added to the hits tested here. Target gene annotations were cross referenced on EcoCyc (41) before testing (**Table S1**). Consistent with our previous data (**Figure 1**), *ΔazoR* was comparable to *wt* (**Figure 2A, Table S1**). A stringent cutoff of 95%confidence interval of all genes tested revealed 22 gene deletions that had significantly impaired activity (**Figure 2A**). The most dramatic loss-of-function was found for *Δfnr* (fumarate and nitrate reduction regulator) resulting in a 2.3-fold decrease in azo dye depletion. We validated the role of *fnr* in FD&C Red No. 40 depletion by constructing a clean deletion (**Figure 2B**) then confirming the deletion by gel electrophoresis (**Figure S1C**) and Sanger sequencing (**Figure S1D**). Activity was restored through complementation of *fnr* on the pCA24N plasmid from the ASKA collection (38) (**Figure 2C**).

**FIG 2.**
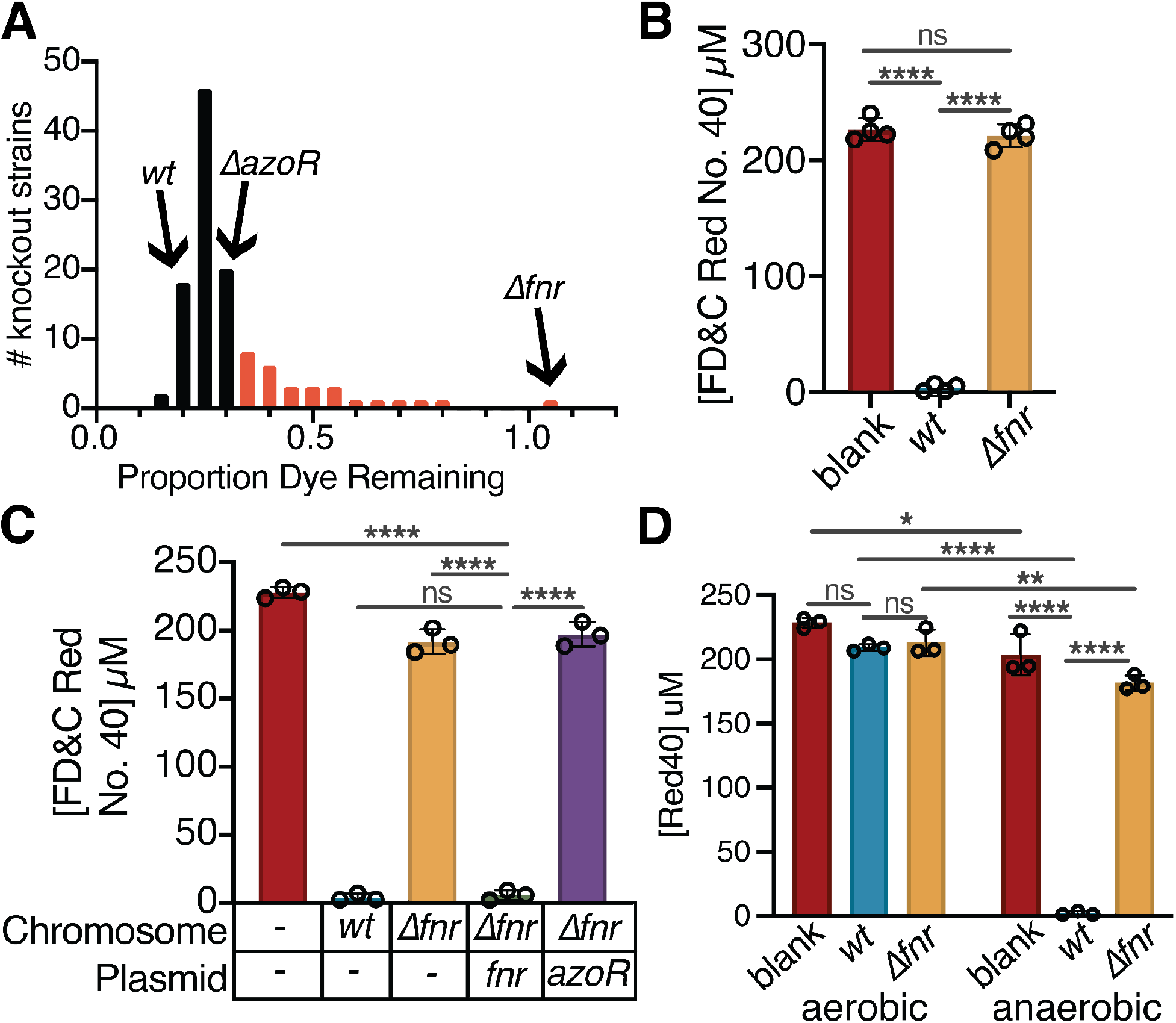
The *fnr* regulator is necessary for azoreductase activity in *E. coli*. (A) A targeted screen of 115 deletions of genes that encode enzymes and regulators revealed 21 loss-of-function strains (red bars, more dye remaining than the 95% confidence interval of the mean; n=6 biological replicates/strain across 2 independent experiments). The *Δfnr* (fumarate and nitrate reduction regulator) strain had the most extreme phenotype (complete loss-of-function). No significant gain-of-function phenotypes were observed. (B) Removal of the kanamycin cassette from Δ*fnr* still leads to a complete loss-of-function (n=4 biological replicates/strain). (C) Complementation of *fnr* but not *azoR* rescues the azoreduction phenotype. Samples all induced with 0.1 mM IPTG at mid-exponential growth phase. (D) *E. coli* azo reductase activity is significantly greater under anaerobic conditions. (A-D) All experiments: 24 hours of growth in LB media, with 4.13 mM L-Cysteine and μM FD&C Red No. 40, under anaerobic conditions with the exception of samples in (D). (A-D) Values are mean±stdev. \**p*<0.05, \*\**p*<0.01, \*\*\**p*<0.001, \*\*\*\**p*<0.0001, (B, C) one-way ANOVA, (D) two-way ANOVA. Supernatant was removed from samples and analyzed for residual dye concentration spectrophotometrically. Concentrations calculated based on a standard curve. Limit of detection = 1.2 μM. (C-D) n=3 biological replicates/strain.

FNR plays a key role in oxygen sensing and the transition to anaerobic metabolism (42, 43). Consistent with this, FD&C Red No. 40 depletion was significantly greater under anaerobic conditions (**Figure 2D**). The requirement for *fnr* was not unique to FD&C Red No. 40; significantly decreased dye depletion was observed across a panel of 7 food dyes and 3 azo bond-containing drugs (**Figure S3**). Overexpression of AzoR was insufficient to rescue Δ*fnr* (**Figure 2C**), suggesting that other aspects of the vast *fnr* regulon (44–47) are responsible for the observed phenotype and/or permeability issues result in inadequate dye reaching AzoR in the cytosol (48).

### The FNR-dependent regulators *cyuR* and *crp* contribute to azoreductase potential

While prior studies have extensively characterized the FNR regulon (43, 44, 46, 49), they were performed in different growth conditions and in the absence of azo dyes, motivating us to revisit these results using paired transcriptomics and proteomics (**Figure 3A**). We grew *wt* and Δ*fnr E. coli* in Luria Broth (LB) media and inoculated the cultures with 250 μM FD&C Red No. 40 at either mid-exponential or stationary growth phases. After a 40-minute incubation period, samples were collected and split for paired transcriptomics (RNA-seq) and proteomics analysis. In total, we generated 12.4±6.2 million high quality reads/sample (RNA-seq; **Table S2**) and 387,908 peptides after matching spectra through the MSGF+ database (50) and adjusting for false discovery rate of 0.01 against the MSGF generated decoy database (proteomics; **Table S3**).

**FIG 3.**
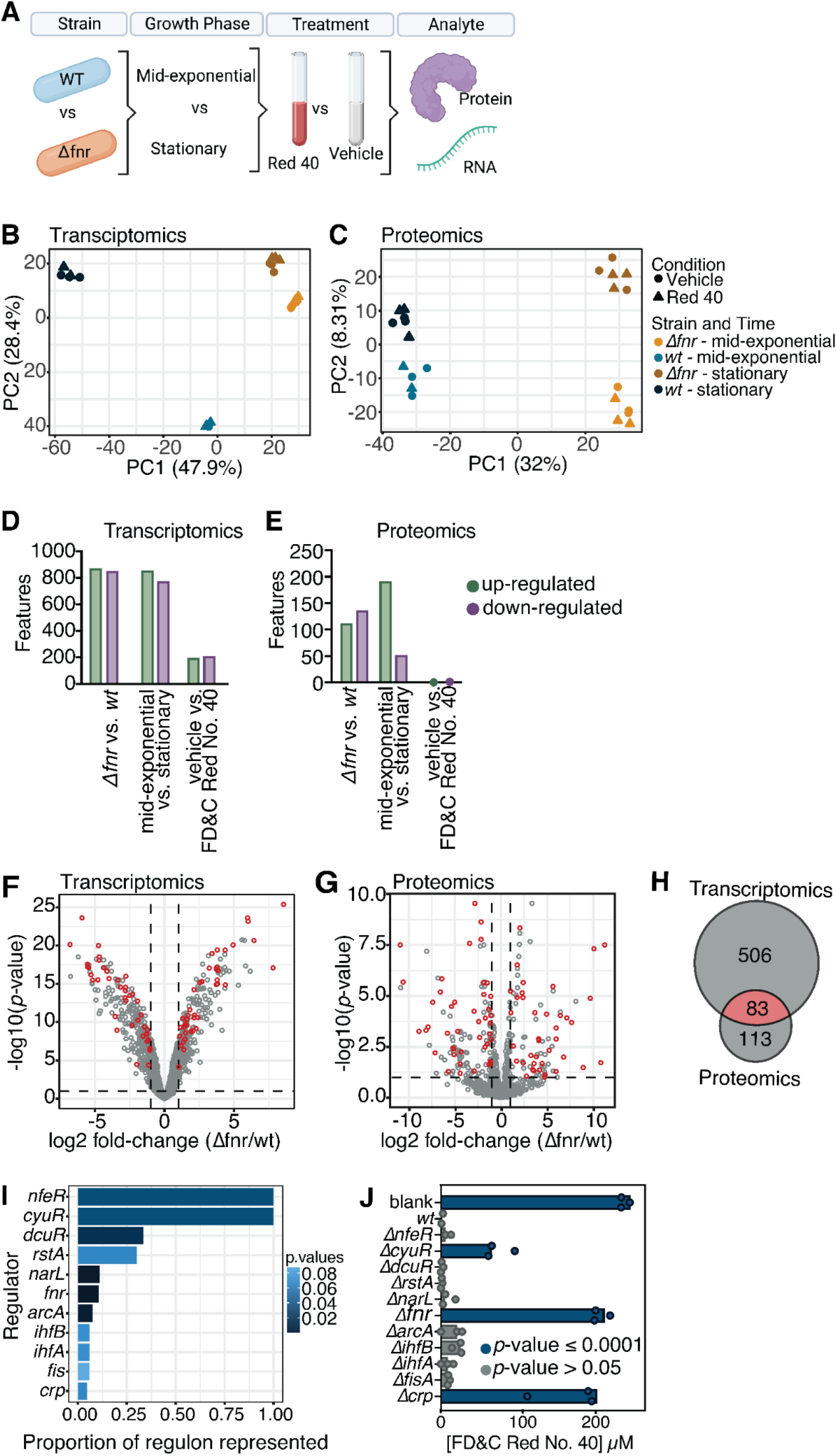
Multi-omic analysis of the *fnr* regulon reveals a complex regulon and the importance of L-Cysteine metabolism. (A) Experimental design. *wt* and *Δfnr* strains were grown to mid exponential or stationary phase, then dosed with 250 μM FD&C Red No. 40 or vehicle control. After 45 mins, cultures were spun down and fractions taken for proteomics or transcriptomics processing. Created with BioRender.com. (B-C) Principal components analysis (PCA) of Euclidean distances for Transcriptomics (B) and Proteomics (C) data sets. Each point represents one sample from the experiment. (D,E) Barplots representing the number of features up-or down-regulated at each contrast of the model used for Transcriptomics (D) and Proteomics (C) data sets. (F,G) Volcano plots for Transcriptomics (F) and Proteomics (G) data sets. Each point represents the average expression of one transcript or protein, respectively. Differential expression is based on the strain level contrast. Significantly differentially expressed features (*p_adj_*<0.1, log_2_ fold-change≥|1|) are indicated by dashed lines, comparing *Δfnr* to *wt*. (H) Venn diagram of the number of differentially expressed features (*p_adj_*<0.1, log_2_fold-change >1) in the two datasets. (I) Regulons enriched in the overlapping features of transcriptomics and proteomics data sets (EcoCyc, *p_adj_*<0.1, BH corrected) of differentially expressed genes (*p_adj_*< 0.1, log_2_ fold-change>|1|, limma). (J) Azo depletion activity by enriched regulator knockout strains (Keio Collection). Values are mean±stdev.

Principal coordinates analysis revealed a clear separation of profiles between strains at the RNA (**Figure 3B**) and protein (**Figure 3C**) levels. As expected (51–53), we also saw a clear difference in expression profile between growth phases; however, the separation between strains was maintained at both timepoints (**Figure 3B,C**). There was a more minimal impact of FD&C Red No. 40 (**Figure 3B,C**).

To identify differentially expressed genes that could account for the observed differences in azo reductase potential, we modeled the complex interaction between growth phase, strain, and treatment (**Figure 3D-E, Table S4**). Specifically, we continued our analyses using the differences found using the strain contrast since so few features were significantly different when taking all conditions contrasts into account. Consistent with the low signal for treatment, the contrast for FD&C Red No. 40 only gave one significantly depleted protein, Flagellar Hook Protein (FlgE). For transcriptomics, we identified 219 significantly upregulated features at FD&C Red No. 40 contrast (*p_adj_*<0.1); however, we decided to move forward with the strain contrast so that transcriptomics and proteomics datasets could be compared. In total, we identified 506 (RNA) and 133 (protein) differentially expressed features between strains (log_2_ fold-change>|1|, *p_adj_*<0.1; **Figure 3F,G**). 83 of these features were consistently differentially expressed in the same direction in both datasets (**Figure 3H**), including 63 annotated enzymes and 11 annotated reductases (**Table S5**). Of note, *azoR* was slightly significantly upregulated in the *wt* strain (**Figure S4A**), with a trend towards increased peptide intensity (**Figure S4B**). As expected, *fnr* transcripts were undetectable in the Δ*fnr* strain (**Figure S4C**).

To further narrow down the subsets of the *fnr* regulon involved in azo reductase activity, we searched for additional regulators in our list of differentially expressed genes. Features that were significantly upregulated in both RNA-seq and proteomics in *wt* compared to Δ*fnr* (log_2_ fold-change> |1|, *p_adj_*<0.1) were analyzed for direct regulators using EcoCyc (see *Methods*). This analysis revealed 11 regulators whose regulons were significantly enriched in our gene set (**Figure 3I**). Reassuringly, *fnr* was the most significantly enriched regulator. We went on to test the azoreduction phenotype of Keio collection knockouts of all 11 regulators (**Figure 3J**). Two regulators that were not in our original screen, *cyuR* (detoxification of L-Cysteine regulator) and *crp* (cAMP-activated global transcriptional regulator), demonstrated a significant loss-of-function compared to *wt* (*p*<0.0001, one-way ANOVA). Crp regulates >100 genes during both aerobic and anaerobic growth (54–57). In contrast, *cyuR* regulates just two genes involved in L-Cysteine metabolism: *cyuA* (L-Cysteine desulfidase) and *cyuP* (L-Cysteine utilization permease) (58–60). Given the tractable size of the *cyuR* regulon and emerging evidence that L-Cysteine metabolism can enable degradation of azo bonds (34, 61, 62), we opted to focus on *cyuA* and *cyuP*.

### FNR-dependent L-Cysteine metabolism is required for degradation of azo bonds

L-Cysteine metabolism by *E. coli* generates hydrogen sulfide (H_2_S) as a dead-end metabolite (**Figure 4A**) (59, 63), which has been previously implicated in azo dye degradation (34, 61, 62). Thus, we hypothesized that *fnr* controls H_2_S production and thus azo bond degradation by increasing the expression of the *cyuR*-regulated *cyuA* and *cyuP*. First, we tested the requirements for L-Cysteine and H_2_S in azoreduction. To maintain anaerobic conditions, we routinely supplement our media with L-Cysteine (64–66). Removal of L-Cysteine impaired the ability of *E. coli* to clear FD&C Red No. 40 (**Figure 4B**). L-Cysteine promoted FD&C Red No. 40 depletion in a dose-dependent manner (**Figure 4C**), suggesting that L-Cysteine is a key substrate for azo reductase activity during *in vitro* growth under anaerobic conditions. Anaerobic conditions in LB media lacking L-Cysteine, equilibrated in the anaerobic chamber for one week were confirmed by eye using the indicator resazurin. Notably, L-Cysteine alone was unable to deplete FD&C Red No. 40 (**Figure 4D**), while addition of sodium sulfide demonstrated a dose-dependent reduction in sterile media (**Figure 4E**). To test the relationship between L-Cysteine and H_2_S, we quantified H_2_S in whole cell culture using the methylene blue assay (see *Methods*). H_2_S levels were below our limit of detection (1.3 μM) in the Δ*fnr* strain, significantly less than the triple-digit μM levels produced by *wt* (**Figure 4F**). We also quantified the amount of FD&C Red No. 40 depleted in these samples and saw the inverse trend (**Figure 4G**).

**Figure 4.**
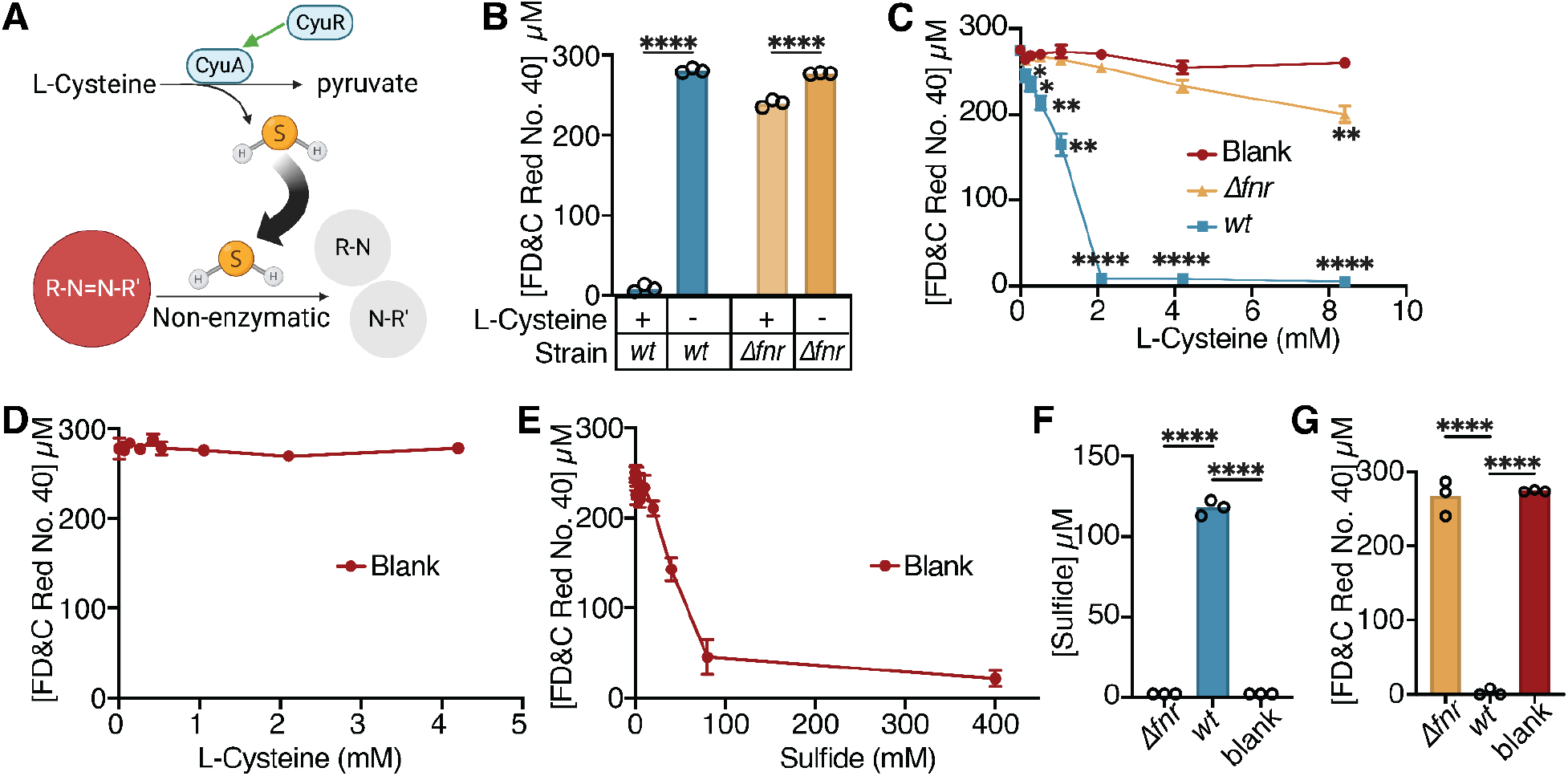
The *fnr* regulator is necessary for the metabolism of L-Cysteine to hydrogen sulfide, enabling the reduction of azo bonds. (A) Schematic depicting L-Cysteine metabolism pathway in *E. coli* which metabolizes L-Cysteine to pyruvate, releasing hydrogen sulfide. Created with BioRender.com. (B) Significant differences in FD&C Red No. 40 dye depletion is observed when *wt E. coli* is grown with or without L-Cysteine in the media. The *Δfnr* strain was also significant at the highest concentration. (C) *wt* and *Δfnr E. coli* was incubated in decreasing concentrations of L-Cysteine. Statistics refer to comparison to blanks in a one-way ANOVA. (D) Increasing L-Cysteine concentrations in LB with 250 uM FD&C Red No. 40. (E) Increasing Sodium Sulfide concentrations in LB with 250 uM FD&C Red No. 40 with 0.05% L-Cysteine. (F) Hydrogen sulfide production by *wt* and *Δfnr E. coli*. (G) FD&C Red No. 40 depletion by *wt* and *Δfnr E. coli*. \**p*<0.05, \*\**p*<0.01, \*\*\**p*<0.001, \*\*\*\**p*<0.0001, two-way ANOVA (B) or one-way ANOVA (C,F,G). All experiments: 24 hours of growth in LB media, with 4.13 mM L-Cysteine and 250 μM FD&C Red No. 40 unless otherwise noted. Supernatant was removed from samples and analyzed for residual dye concentration spectrophotometrically. Concentrations calculated based on a standard curve. Limit of detection = 1.2 μM (n=3 biological replicates).

Next, we investigated if H_2_S alone was the *fnr*-driven azoreductase mechanism for FD&C Red No. 40 depletion. We utilized single and multi-gene knockouts for the *cyuR* regulon. As previously mentioned, *cyuR* regulates both the L-Cysteine transporter, *cyuP* and L-Cysteine desulfurase, *cyuA* (59) (**Figure 5A**). Consistent with our hypothesis, both *cyuA* and *cyuP* were significantly decreased at the transcript level in Δ*fnr E. coli* relative to *wt* controls (**Figure 5B**)*. cyuR* was also significantly increased at the transcript level (**Figure S4D**). The *cyuP* and *cyuR* knockout strains produced significantly less sulfide (**Figure 5C**) and depleted significantly less FD&C Red No. 40 (**Figure 5D**) in L-Cysteine-containing media. The combined deletion of *cyuR* and *azoR* did not significantly impact sulfide levels (**Figure 5E**); however, there was a slight but statistical significance impairment in azo dye depletion (**Figure 5F**).

**FIG 5.**
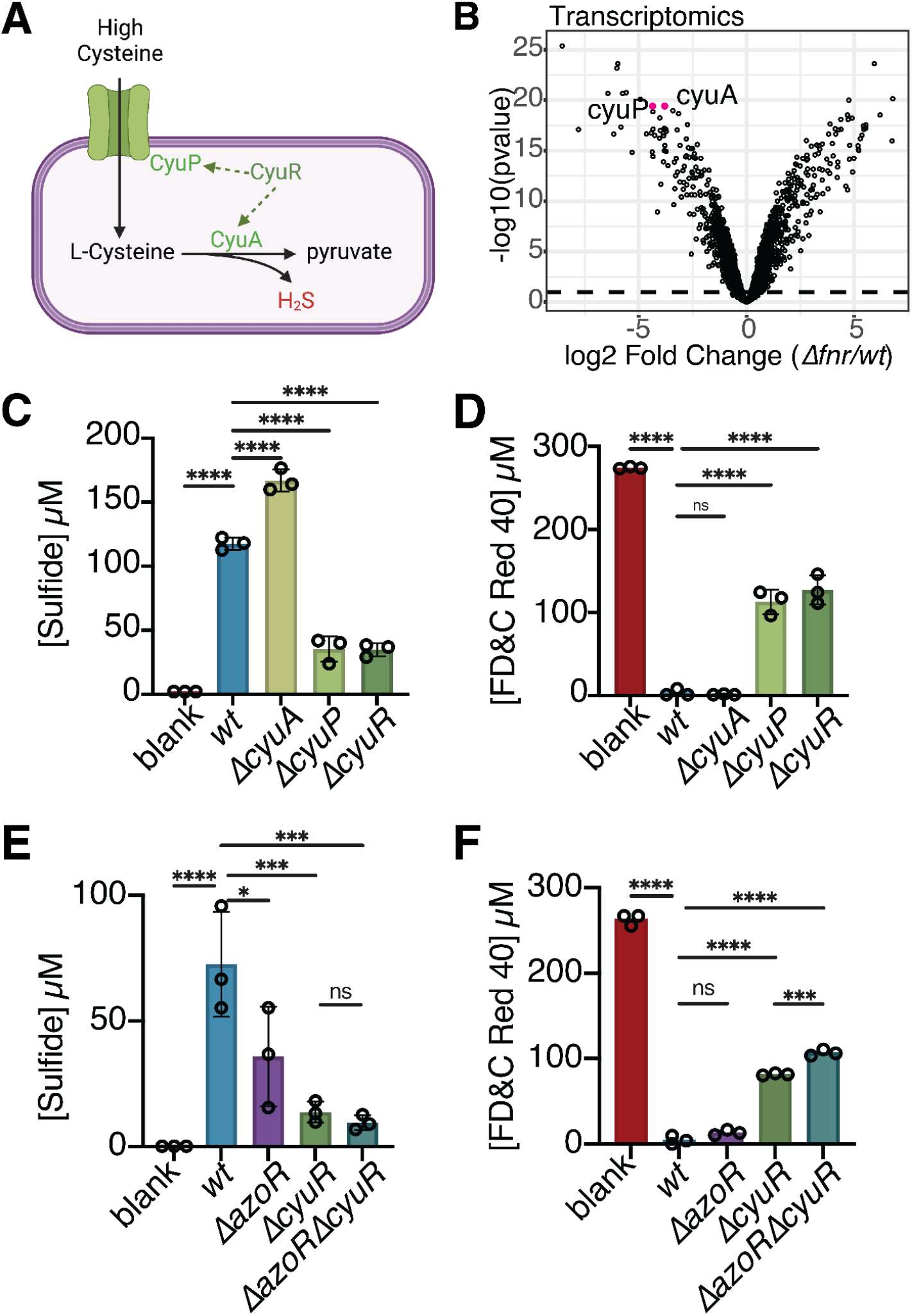
L-Cysteine import and metabolism genes impact bacterially produced sulfide and FD&C Red No. 40 depletion. (A) Model of the *cyuR* regulon’s role in L-Cysteine metabolism. CyuP acts as a L-Cysteine transporter into the cell, while CyuA is a L-Cysteine desulfidase. Created with BioRender.com. (B) Volcano plot for transcriptomics data: *cyuA* and *cyuP* are highlighted in pink. Each point represents the average expression of one transcript. Differential expression is based on the strain level contrast. (C-D) Paired sulfide (C) and FD&C Red No. 40 (D) were measured from *cyuR* regulon knockouts and *cyuR* itself (Keio collection). (E-F) Paired sulfide (E) and FD&C Red No. 40 (F) were measured from *ΔazoR, ΔcyuR*, and *ΔazoRΔcyuR* strains. Values are mean±stdev. \**p*<0.05, \*\**p*<0.01, \*\*\**p*<0.001, \*\*\*\**p*<0.0001, one-way ANOVA. (C-D) n=3 replicates/strain.

## DISCUSSION

Our results provide mechanistic insight into the degradation of azo dyes by human gut bacteria. Surprisingly, the canonical azoreductase enzyme of *E. coli* is dispensable for whole cell azoreductase activity. Instead, the shift to anaerobic growth is critical. We dissected a pathway through which oxygen is sensed by FNR, leading to upregulation of the *cyu* operon, increased uptake of L-Cysteine, enhanced production of H_2_S, and non-enzymatic azo bond reduction (**Figure 6**).

**FIG 6.**
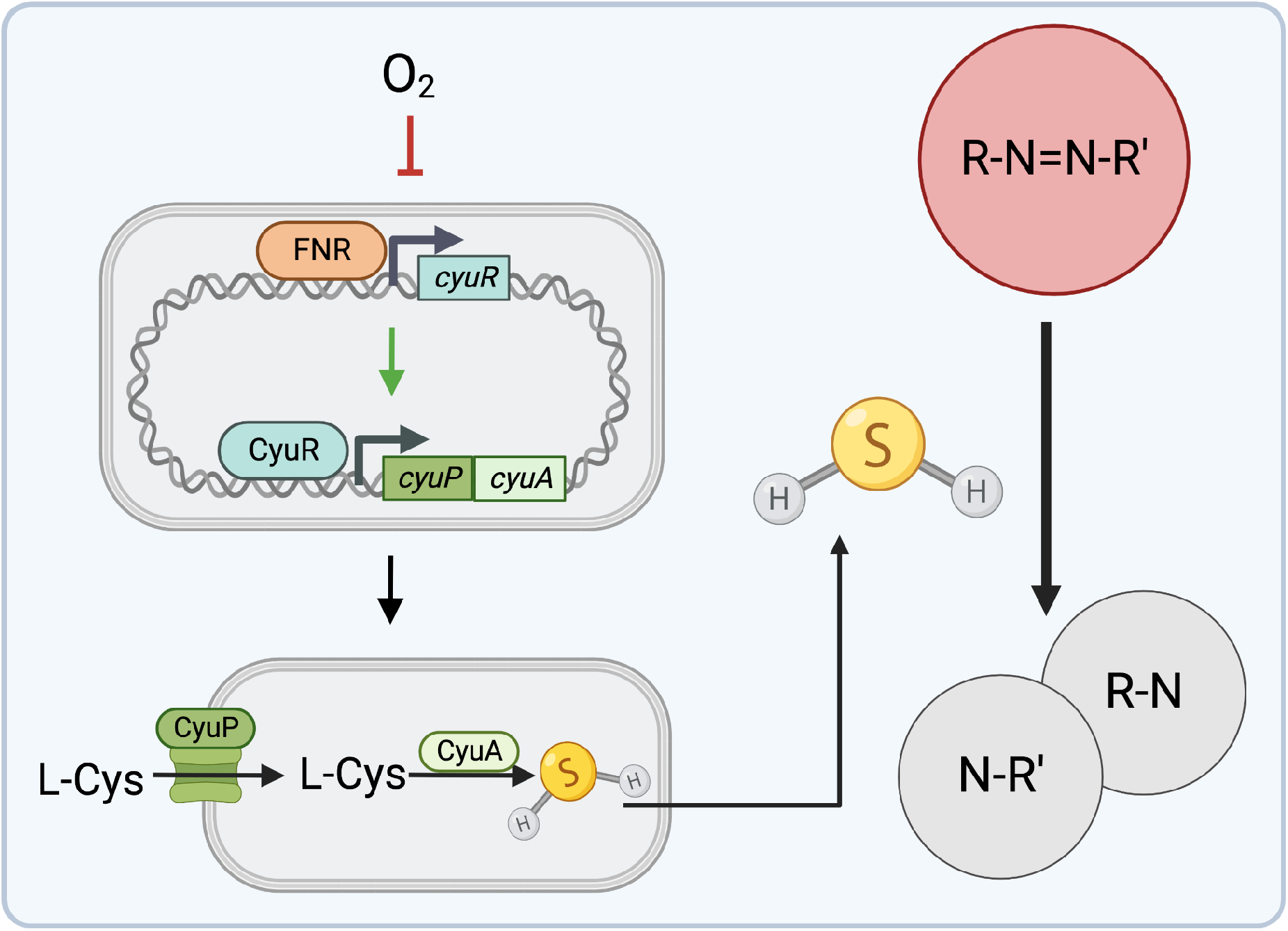
Working model for how the host environment shapes gut bacterial azo reduction. Under anaerobic conditions, FNR activates the transcription of *cyuR*. CyuR in turn activates expression of the *cyuAP* operon. CyuP transports L-Cysteine into the cell, where it is metabolized to H_2_S through the CyuA desulfidase. Released H_2_S can reduce the azo bond of azo compounds. Created with BioRender.com.

These findings provide a cautionary tale for the ability to extrapolate results from biochemical studies of purified proteins to infer the metabolic activity of cells and complex microbial communities. Despite a rich literature on *E. coli* AzoR *(24, 48, 67)*, we found that the *azoR* gene is dispensable for azo reduction by *E. coli* during anaerobic growth. These findings emphasize the importance of pairing biochemical and heterologous expression studies of enzyme function with methods for gene deletion in the original strain. Future work should extend this approach to the other bacterial species with biochemically characterized azoreductase enzymes (25, 29, 68, 69). For species where genetic tools do not exist (the vast majority of human gut bacteria), one could leverage natural strain-level variation to test the association between gene presence and sequence variation with metabolic activity (37).

We found that the global regulator FNR is essential for *E. coli* azoreduction, highlighting the importance of environmental oxygen for the gut bacterial metabolism of drugs and excipients. Confirming the prior work on FNR in different growth conditions (42, 57), we found that the FNR regulon is vast, including hundreds of genes involved in different aspects of anaerobic metabolism. Interestingly, efforts to utilize bacterial azoreduction to degrade dye runoff in water systems also require anaerobic conditions, suggesting that our findings may be generalizable to microbial communities outside of the human gut (23, 70, 71). Importantly, *E. coli* exhibits features of both aerobic and anaerobic growth within the GI tract (57, 72), depending upon its physical niche and host pathophysiology (73, 74). Our results suggest that inter-individual variations in the environment in which *E. coli* grows could have downstream consequences for its ability to act upon azo bond containing compounds. More work is necessary to assess how the redox potential of the gut lumen and other environmental factors shape gut bacterial azo reduction in the context of host health and disease.

Surprisingly, we found that FNR controls azo reductase potential through the reduction of azo bonds by H_2_S. Given that L-Cysteine is the major source of H_2_S for *E. coli*, these results provide an additional layer of environmental control for azoreduction. Differences in dietary protein, host proteoglycans, and/or L-Cysteine biosynthesis by other members of the gut microbiota could all potentially influence the activity of *E. coli* and its ability to act on azo bonds. Future studies leveraging gnotobiotic mice, defined dietary interventions, and isotope tracing could be informative in this regard.

Our results indicate that the L-Cysteine utilization regulator *cyuR*, among others, is under FNR control, emphasizing the complex multi-level regulatory network contributing to bacterial azoreduction. The *cyu* operon is the main source for L-Cysteine import and metabolism by *E. coli* (59, 75). We also found that H_2_S production requires *cyuR*. Interestingly, we found that the L-Cysteine uptake transporter, CyuP, was more important than the enzyme CyuA, suggesting that the import of L-Cysteine could provide substrate for additional pathways for H_2_S production. CyuA only functions anaerobically (59), consistent with our observations that azoreduction only occurs anaerobically and is regulated by FNR and CyuR. Moreover, the key role of L-Cysteine import suggests that amino acid availability from diet and/or host tissues could be critical for gut bacterial azoreduction. Future studies with mouse models and various dietary amino acid interventions could provide an opportunity to modulate microbiome azoreductase activity.

Deletion of both *cyuR* and *azoR* did not fully abolish *E. coli* azoreduction. Thus, additional pathways for L-Cysteine metabolism and/or as-of-yet undiscovered enzyme(s) may contribute to the metabolism of azo dyes by *E. coli*. In our initial screen, we found 22 deletion strains with a partial loss-of-function phenotype, all of which encode enzymes theoretically capable of azo reduction. Of these, 4 were also identified in our paired proteomic and transcriptomic dataset (**Table S6**), providing viable candidates for further biochemical characterization and/or complementation screens in the Δ*fnr* background.

H_2_S production is widespread throughout gut bacterial species (76), suggesting that far more bacterial taxa are capable of azo reduction than previously appreciated. L-Cysteine can be made from a variety of different sources, thus methionine and other sulfur starting points could impact the gut microbiota’s H_2_S production capacity. Multiple *Desulfovibrio* species found in the GI tract can reduce sulfate to H_2_S (77–80). Thus, GI levels of H_2_S represent the net effect of multiple distinct pathways, which will be important to consider when translating these results into humans. As a first step, studies in gnotobiotic mice with defined microbial communities representing distinct assemblages of H_2_S producers would help to better understand the relative importance of each species for azo dye degradation under different dietary and host selection pressures.

There are multiple limitations of this study. We focused entirely on the model gut Proteobacterium *E. coli* due to its genetic tractability and long history of molecular research; however, the generalizability of the pathway we described to other gut bacteria remains unclear. The development of genetic tools that permit similar mechanistic dissections into these more exotic gut bacteria will be essential for studying the conservation of enzymatic and non-enzymatic azo bond degradation in diverse gut bacterial species. The physiological relevance of our findings remains unclear, requiring more work in mice or other preclinical models. It will also be important to test the clinical relevance of these findings in the context of diseases intentionally treated with azo bond containing drugs (*e.g*., inflammatory bowel disease) or diseases like cancer that could potentially be exacerbated by dietary exposures to azo dyes. Our results emphasize that in addition to metagenomics, it will be critical to measure key environmental factors in human clinical cohorts like oxygen and amino acid levels.

Despite these limitations, our results clearly emphasize the importance of environmental factors in controlling a clinically relevant metabolic activity of the human gut microbiome. These experiments are particularly timely given the rapid increase in food dye intake due to their incorporation in processed foods (81) and emerging evidence that they have unintended consequences for gut epithelial (18) and immune (22) cells. Continued progress towards dissecting the mechanisms responsible for this metabolic activity in *E. coli* and other experimentally tractable gut bacterial species is a key step towards predicting and controlling the location and extent of azo dye metabolism.

## MATERIALS AND METHODS

### Bacterial strains, media, and chemicals

Bacterial strains used in this study are listed in **Table S7**. *E. coli* was grown in Lysogeny broth (LB) supplemented with 0.05% L-Cysteine under anaerobic conditions at 37°C unless otherwise noted. L-Cysteine was purchased from Sigma-Aldrich (St. Louis, MO).

### Assaying whole cell *E. coli* depletion of azo compounds

Cultures were grown in Luria Broth (LB) medium supplemented with 250 μM of azo dye and 0.05% L-Cysteine. Azo dye stocks were prepared in DMSO at a concentration of 25 mM and used at a final concentration of 1% (v/v). Cultures were incubated at 37°C. All incubations were performed in a COY Laboratory Products Inc anaerobic chamber (Grass Lake, MI) with the following atmosphere: 10% H_2_, 5% CO_2_, and 85% N_2_, All reagents were equilibrated in the anaerobic chamber for at least 24 hours before use. After 24 hours of growth, samples were removed, spun down to remove cells, and analyzed for residual dye concentration by measuring absorbance at appropriate wavelengths. Wavelengths used: 450 nm (FD&C Red No. 40, FD&C Yellow No. 6, D&C Orange No. 4), 360 nm (Balsalazide, Sulfasalazine, Olsalazine, Phenazopyridine), 420 nm (D&C Brown No. 1), 500 nm (D&C Red No. 6, FD&C Red No. 4), 535 nm (D&C Red No. 33). Residual dye concentration was determined by creating a standard curve of dye concentrations in LB and measuring absorbance at appropriate wavelengths. Concentration was calculated using the equation of the fitted standard curve line.

### Clean deletion creation from Keio collection strains

Competent cells were created from cultures of the Keio collection knockouts for *azoR* and *fnr* knockouts. Cells were electroporated with vector pSIJ8 (82) and transformants were selected for on LB supplemented with 25 μg/ml ampicillin at 30°C. Removal of the kanamycin cassette at FRT sites was done on liquid culture of transformant using 50 mM L-rhamnose for 4 hours, then patch streaked to identify loss of kanamycin resistance. Transformants were streak-purified two times, then PCR was performed for confirmation of kanamycin cassette removal. Strains were cured of plasmid at 37°C and then patch-streaked to confirm loss of ampicillin resistance.

### Purified enzyme preparation

The ASKA collection (38) strain expressing *azoR* was streaked from glycerol stock on LB with 30 μg/ml chloramphenicol. A single colony was used to inoculate an overnight culture of 25 ml of LB with 30 μg/ml chloramphenicol aerobically at 37°C. The overnight was used to inoculate 1 L of LB with 30 μg/ml chloramphenicol at a 1:10 dilution factor. Sample was grown with shaking aerobically to an optical density of 0.5 at 600 nm. Expression of *azoR* was induced with Isopropyl β-d-1-thiogalactopyranoside (IPTG) at 1 mM concentration for 3 hours. Cells were then cooled on ice water for 10 mins and centrifuged at 3,500 x g for 15 mins, washed with ice cold PBS, spun again, then cell pellets were frozen overnight at −20°C. Pellets were thawed for on ice, then resuspended in Buffer A (50 mM HEPES, 300 mM NaCl, 10 mM imidazole, pH 7.5) and sonicated with the following program: 5 mins, 4 sec on, 4 sec off, at 30% amplitude. Protease tablets (Sigma Aldrich) were crushed in buffer A and added to the sonicated lysate. Lysate was centrifuged for 20 mins at 16,000 RPM. Supernatant was applied to a 5 ml nickel-nitrilotriacetic acid column (Qiagen, Valencia, CA) that had been equilibrated with buffer A at a rate of 1 ml/min. The column was washed with 5 column volumes of buffer A, then 5 column volumes of buffer B (50 mM HEPES, 300 mM NaCl, 20 mM imidazole, pH 7.5). Protein was eluted with Buffer C (50 mM Hepes, 300 mM NaCl, 300 mM imidazole) and collected in 5 ml portions. Fractions were analyzed for protein content using a sodium dodecyl sulfate-polyacrylamide gel electrophoresis (SDS-PAGE) gel. Fractions containing proteins corresponding to AzoR molecular weight were pooled and dialyzed with 3 L of buffer D (50 mM Hepes, 300 mM NaCl, 15% glycerol by weight, in cold water, pH 7.5) overnight at 4°C. Protein concentration was determined using Nanodrop (Thermo Scientific).

### Purified Enzyme Experiments

AzoR reactions with azo dyes were carried out in 20 mM sodium phosphate buffer, 100 μM azo dye (unless otherwise noted), 2000 μM NADH, and 20 μM FMN with varying concentrations of enzyme as appropriate. Reactions were activated by the addition of NADH. Dye concentration was monitored at absorbance 450 nm every 30 seconds after a linear shake of 15 seconds. All reactions were performed aerobically at 37°C.

### Keio collection screening for alternative azo reducers

Homologs to genes implicated in azoreduction by *Shewanella oneidensis* MR-1 (39) were identified by first gene name and second gene function (tested or predicted) in *E. coli*. Other stains of interest were identified based on gene name and function. All strains were cross referenced on Ecocyc (41). Strains were revived from glycerol stocks on LB agar with 25 μg/ml kanamycin. Single isolates were picked from agar plates and grown overnight in 1 ml of LB supplemented with 0.05% L-Cysteine in a 96-deep well plate under anaerobic conditions (see above). For azoreduction assay, previously anaerobically equilibrated liquid media was supplemented with 25 μM FD&C Red No. 40 and 0.05% L-Cysteine. 1 ml of media was aliquoted to 2 ml deep 96 well plates and strains were inoculated at 1:100 from overnight cultures in triplicate. Plates were sealed with TempPlate Sealing Foil (USA Scientific, Cat #2923-0110) and allowed to grow for 24 hours. After 24 hours, cells were spun down and 100 μl of supernatant was used to measure dye absorbance at 450 nm. Proportion dye remaining was calculated by dividing the absorbance of each well by the average absorbance of uninoculated control wells. Where standard curves were used, a 1:2 dilution series of FD&C Red No. 40 was made in LB with 0.05% L-Cysteine in triplicate and dye absorbance measured at 450 nm. Concentrations were calculated using the equation of the standard curve line in Excel.

### Validation of Keio collection knockouts

To ensure the knockouts used from the Keio collection were indeed the annotated gene, primers were designed up and downstream of the gene of interest. PCR and Sanger sequencing of the kanamycin insert and surrounding regions were performed to ensure the expected band size and gene alignment. Primers are listed in **Table S8**.

### Overexpression strain construction and assay

Overexpression strains were made by preparing competent cells for the background strain of interest. Plasmids were prepared by growing up overnight cultures from the ASKA strain of interest, then using Qiagen Plasmid Mini Kit (Qiagen; Cat #12125) kit to extract the pCA24N plasmid with the gene of interest. The plasmid preparation was desalted using a membrane filter on water, then electroperated to the background strain of interest. Correct insertion was ensured by growth of the transformant on chloramphenicol. A miniprep was also made from the transformant and then PCR and Sanger sequencing done to ensure the expected gene insertion.

### RNA-Sequencing and proteomics sample growth and exposure

100 ml of overnight cultures of both *wt* and Δ*fnr E. coli* were grown in LB supplemented with 0.05% L-Cysteine under anaerobic conditions. Overnight culture OD_600 nm_ was measured for both cultures and normalized to a starting OD_600 nm_ of 0.05 in 100ml of LB with 0.05% L-Cysteine in 12 replicates for each strain. OD_600 nm_ was monitored until mid-exponential phase for each strain (previously determined). At mid-exponential 3 replicates for each strain were treated with a final concentration of 250 μM FD&C Red No. 40 in DMSO, or 1% DMSO. After 45 mins of exposure, 15 ml of the sample was removed for RNA extraction, while the remaining volume was used for proteomics. Both volumes samples were spun down to pellet cells, supernatant removed, and cell pellets were flash frozen in liquid nitrogen then stored at −80°C. The remaining samples were allowed to reach stationary phase, then they were treated with a final concentration of 250 μM FD&C Red No. 40 in DMSO, or 1% DMSO. Samples were processed the same way as for mid-exponential.

### RNA-sequencing sample preparation

1 μl of TRI Reagent (Sigma Aldrich catalog number T9424) was added to bacterial pellets and incubated for 10 mins, samples were then transferred to 2 ml Lysing Matrix E tubes (MP Biomecials, catalog number 116914050). Cells were lysed for 5 mins in the bead beater at room temperature. Next, 200 ul of chloroform was added. Samples were vortexed for 15 sec and incubated at room temperature for 10 mins. Samples were then centrifuged at 16,000 x g for 15 mins at 4°C. 500 μl of the upper aqueous phase was transferred to a new tube, 500 μl of 100% ethanol added, and vortexed to mix. Mixture was transferred to a spin column (PureLink RNA Mini Kit; Life Technologies; catalog number 12183025) and centrifuged at ≥12,000 x g for 30 sec, discarding flow-through, until all of the material had been added to the column. To the spin column, 350 μl Wash Buffer I (PureLink RNA Mini Kit; Life Technologies; catalog number 12183025) was added, then the column was centrifuged at ≥ 12,000xg for 30 sec, discarding flow-through. 80 μl of PureLink DNase mix was added to the column and incubated at room temperature for 15 mins. Next, 350 μl of Wash Buffer I (Purelink RNA mini kit) was added and the column spun at >12000 x g for 30 sec. The column was transferred to a new collection tube, and 500 μl Wash Buffer II was added, followed by centrifugation at ≥12,000 x g for 30sec, discarding flow-through. The column was centrifuged at ≥12,000 x g for 60 sec and dried, then moved to a collection tube. Then, 50 μl RNase-free water was added, and the column was incubated at room temperature for 1 min. Finally, the column was centrifuged for 1 min at ≥12,000xg, retaining the flow-through, which contained total RNA.

Next, samples were DNase treated again using TURBO-DNase (Ambion; ThermoFisher catalog number AM2238) and incubated at 37°C for 30 mins. Next, RNA Ampure XP Beads were used to clean up the reactions. 1.8 volumes of the RNA Ampure XP beads were added to 1 volume of RNA sample and allowed to sit for 5 mins at room temperature. Tubes were placed on a magnetic stand until the liquid cleared. Then liquid was removed and beads washed with 200 μl of 100% Ethanol. Samples were incubated for 30 sec, then removed. This was repeated twice more. Then, samples were allowed to dry for 5 mins, removed from the magnetic stand, and 30 μl of RNase-free water was added. This was incubated for 2 min at room temperature then placed back on the magnetic rack where liquid was collected after turning clear.

rRNA was removed from total RNA using Ribominus Transcriptome Isolation Kit for Bacteria and Yeast (Invitrogen; catalog number K155004, LOT: 2116711), following manufacturer’s protocol. RNA fragmentation, cDNA synthesis, and library preparation were performed using the NEBNext Ultra RNA library Prep Kit for Illumina and NEBNext Multiplex Oligos for Illumina (Dual Index Primers) (Ipswich, MA). Samples were dual-end sequenced (2 x 75 bp) using the NextSeq Mid Output platform (**Table S2**). Reads were mapped to the *E. coli* K12 BW25113 genome sequence (NCBI Reference Sequence: GCA_000750555.1) using Bowtie2 (83) and HTSeq (84) was used to count the number of reads to *E. coli* genes. Differential gene expression was analyzed using limma (85). Differentially expressed genes were defined as transcripts exhibiting an absolute log_2_ fold-change ≥1 and a FDR<0.1.

### Proteomics sample preparation

Cell pellets were washed three times with 5 ml of cold PBS and resuspended in 1 ml, transferred to 2 ml Lysing Matrix E tubes (MP Biomecials, catalog number 116914050) and lysed via bead beating for 1 min, then resting tubes on ice for 2 mins. This was repeated twice. Samples were centrifuged for 15 min at 4°C at 16,000 x g. Next, a BCA assay was performed to quantify protein concentration in the lysate supernatant. Next, urea was added to the samples to a target concentration of 8 M followed by an appropriate volume of Dithiothreitol to obtain a 5 mM concentration. Samples were then incubated at 60°C for 30 mins. While samples were incubating, the trypsin was pre-activated for 10 mins at 37°C. Next, samples were diluted 10-fold with 100 mM NH_4_HCO_3_. Next, 1 M CaCl_2_ was added at an appropriate volume to create a final sample concentration of 1 mM CaCl≤. Samples were then digested for 3 hours with Trypsin at 37°C at a concentration of 1 μg trypsin/ 50 μg protein. Samples were snap frozen.

Next, samples were cleaned with a C_18_ column on a vacuum manifold. The column was conditioned with 3 ml of Methanol, then rinsed with 2 ml of 0.1% TFA acidified water. Sample was run though column, then column was rinsed with 4 ml of 95:5 H_2_O:acetonitrile (ACN), 0.1% trifluoroacetic acid (TFA). Column was allowed to go to dryness, then eluted slowly to dryness with 1 ml of 80:20 ACN:H_2_O, 0.1% TFA into a collection tube. Samples were concentrated in the speed-vac to a volume of approxoimatly 50-100 μl. Protein quantification was done using a BCA test, then samples were stored at −80°C until analysis.

### Proteomics

A Waters nano-Acquity M-Class dual pumping UPLC system (Milford, MA) was configured for on-line trapping of a 5 μL injection at 5 μL/min with reverse-flow elution into the analytical column at 300 nL/min. The trapping column was packed in-house (PNNL) using 360 μm o.d. fused silica (Polymicro Technologies Inc., Phoenix, AZ) with 5 mm Kasil frits for media retention and contained Jupiter C_18_ media (Phenomenex, Torrence, CA) in 5 μm particle size for the trapping column (150 μm i.d.×4cm long) and 3 μm particle size for the analytical column (75 μm i.d.×70cm long). Mobile phases consisted of (A) water with 0.1% formic acid, and (B) acetonitrile with 0.1% formic acid. The following gradient profile was performed (min, %B): 0, 1; 2, 8; 25, 12; 85, 35; 105, 55; 110, 95; 115, 95; 117, 50; 119, 95; 121, 95; 123, 1.

MS analysis was performed using a Velos Orbitrap Elite mass spectrometer (Thermo Scientific, San Jose, CA) outfitted with an in-house made nano-electrospray ionization interface. Electrospray emitters were prepared using 150 μm o.d. ×20 μm i.d. chemically etched fused silica (86). The ion transfer tube temperature and spray voltage were 325 °C and 2.2 kV, respectively. Data were collected for 100 min following a 20 min delay from sample injection. FT-MS spectra were acquired from 400 to 2000 *m/z* at a resolution of 35k (AGC target 3e6) and while the top 12 FT-HCD-MS/MS spectra were acquired in data dependent mode with an isolation window of 2.0 *m/z* and at a resolution of 17.5k (AGC target 1e5) using a normalized collision energy of 30 sec and a 30 sec exclusion time. Extracted MS data were run through the MSGF+ database and filtered by an FDR of >0.01 using the MSGF+ generated decoy database. Differential peptide expression was analyzed using DEP (87) and limma (85). Differentially expressed genes were defined as transcripts exhibiting an absolute log_2_ fold-change ≥1 and a FDR<0.1.

### Hydrogen sulfide quantification

H_2_S was measured using the Cline reaction (88) with modifications. Specifically, samples were put into a zinc acetate buffer (16.7 mM) on ice at a ratio of 1:3 sample:buffer. Then 180 μl of zinc acetate solution was transferred and mixed with 20 μl of cline reagent (2 g of N,N-dimethyl-p-phenylenediamine sulfate with 3 g of FeCl_3_ in 500 ml of cold 50% v/v HCl). The reaction was allowed to proceed for 20-30 minutes at room temperature in the dark. Then absorbances were measured at 670 nm. A standard curve of sodium sulfide was created in zinc acetate and reacted with cline reagents for sulfide concentration calculations.

### Double gene knockout creation

P1 lysates were generated of each strain of interest carrying the kanamycin resistance cassette (Keio collection) adapting methods from previously described techniques (89). Briefly, 150 μl of overnight culture in LB supplemented with 12.5 μg/ml kanamycin was mixed with 1 to 25 μl of P1 phage (previously propagated from ATCC on MG1655). This mixture was incubated for 10 min at 37°C to aid absorption, added to 3 ml of 0.7% agar, and overlaid on pre-warmed LB agar supplemented with 25 μg/ml kanamycin and 10 mM MgSO4. Plates were incubated overnight at 37°C and phage were harvested by adding 5 ml of SM buffer, incubating at room temperature for 10 mins, and breaking the top agar for phage harvest. The mixture was briefly centrifuged to pellet agar, then supernatant was passed through a 100 μm cell straining, then 0.45 μm syringe filter. Lysates were stored at 4°C.

A clean deletion of the recipient strain of interest was created using the above described method with pSIJ8 (82). To transduce the clean recipient strain, 1 ml of an overnight culture of recipient strain was pelleted and resuspended in ⅓ volume of LB 10 mM MgSO4, 5 mM CaCl_2_. 100 μl of cells were mixed with 1 μl to 10 μl of P1 lysate and incubated for 60 min at 37°C. Next, 200 μl of 1M sodium citrate was added with 1 ml of LB to minimize secondary infection. This mixture was incubated at 37°C for 2 hrs, then plated on LB supplemented with 10 mM sodium citrate and 25 μg/ml kanamycin to select for transductants. Transduction was confirmed using PCR and Sanger sequencing. Azoreduction and sulfide activity were tested as described above.

### Statistical analysis

For multiple comparisons of data, an ordinary one-way analysis of variance (ANOVA) test with Dunnet’s correction was chosen, unless otherwise specified. For the targeted knockout screen, a 95% confidence interval of averages from two experiments was calculated using GraphPad Prism 9. Michaelis-menten curves were created in Prism 9. For proteomics and transcriptomics analysis, R package limma was used to determine the fold-change and adjusted *p*-values. Feature count differences (**Figure S4**) and Wilcoxon rank-sum significance testing were done in R. For the regulator enrichment analysis, Ecocyc’s enrichment analysis for “Genes enriched for transcriptional/translational regulators (direct only)” was used with a fisher-exact test with Benjamini-Hochberg Correction.

## Data and material availability

RNA-seq data are available through the NCBI Gene Expression Omnibus (GEO) online data repository under accession number TBD. Proteomics data is available through ProteomeXchange.org, accession number TBD. All other data are provided in the supplemental tables or available upon request.

## SUPPLEMENTAL MATERIAL

Tables S1-S8, XLSX file

## ACKNOWLEDGEMENTS

Funding was provided by the National Institutes of Health (R01HL122593, R01AT011117, R01DK114034, P.J.T.; T32AI0605357, L.P.; ES029319, ES030220, A.T.W.) and the National Sciences Foundation (2018257103, L.P.). P.J.T. is a Chan Zuckerberg Biohub Investigator and held an Investigators in the Pathogenesis of Infectious Disease Award from the Burroughs Wellcome Fund. PNNL is operated by Battelle for the DOE under contract DE-AC06-76RL01830.

Conceptualization, L.P. and P.J.T.; Designed Experiments, L.P., P.S., R.V., P.J.T.; Performed experiments, L.P., R.V. C.M., P.S.; Visualization, L.P.; Supervision, A.T.W., P.J.T.; Writing-original draft, L.P.; Writing-review & editing, P.S., A.T.W., P.J.T.

We thank the Libusha Kelly and Aaron Wright labs for technical assistance with mouse experiments and global proteomics.

P.J.T. is on the scientific advisory boards for Pendulum, Seed, and SNIPRbiome; there is no direct overlap between the current study and these consulting duties. A.T.W. is on the scientific advisory board for Enzymetrics Biosciences; there is no direct overlap between the current study and this role. The other authors declare no competing interests.

## SUPPLEMENTAL FIGURES AND LEGENDS

**FIG S1.**
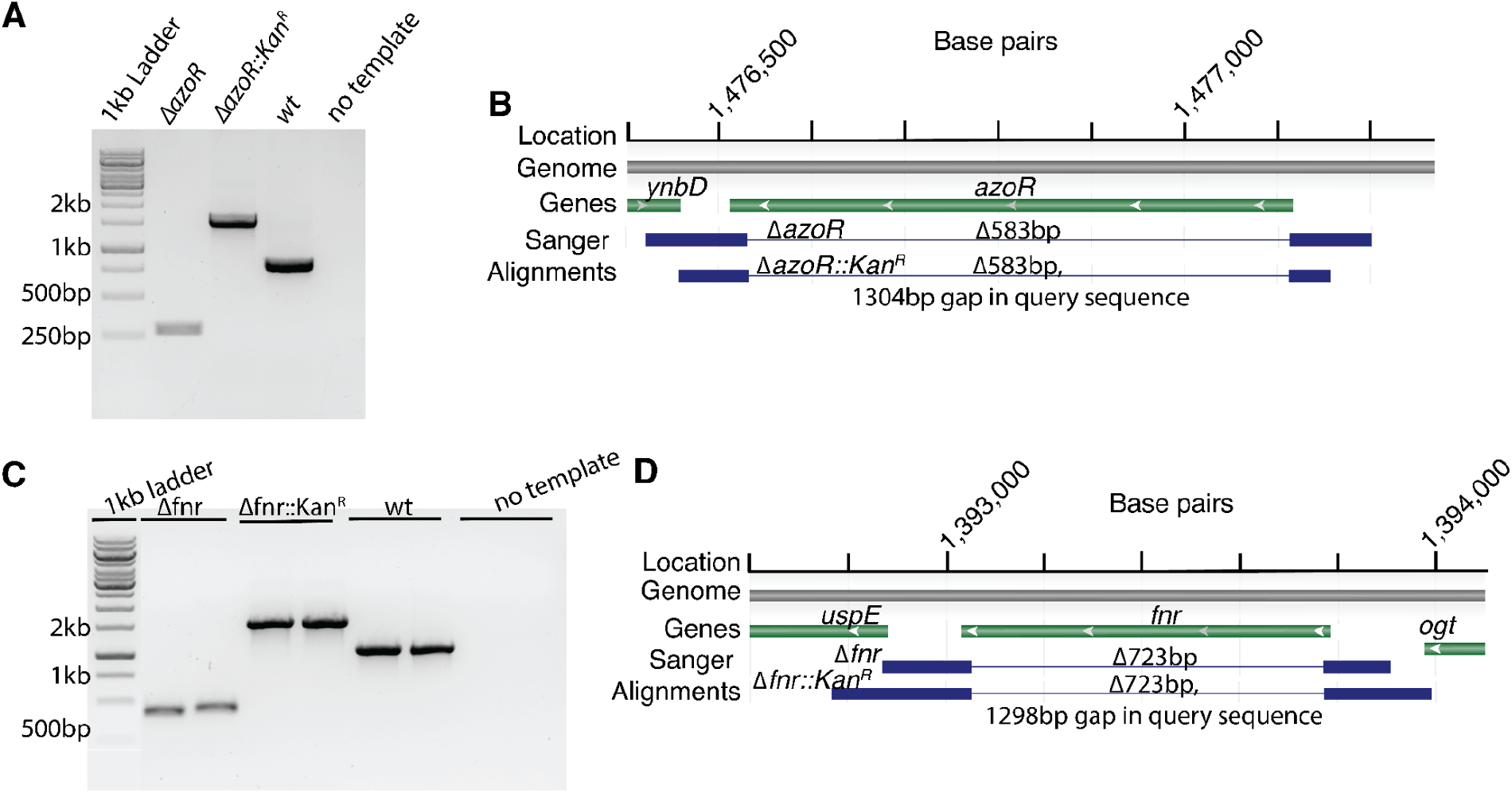
Validation of bacterial strains. (A) Gel electrophoresis of PCR product from *Δ*azoR strains used: *ΔazoR* and *ΔazoR::kanR* compared to wild-type (*wt*). (B) Alignment of the *ΔazoR* and *ΔazoR::kanR* Sanger sequencing results to the *E. coli* K12 BW21153 genome. (C) Gel electrophoresis of PCR product from *Δfnr* strains used: *Δfnr* and *Δfnr::kanR* compared to wild-type (*wt*). (D Alignment of the *Δfnr* and *Δfnr::kanR* Sanger sequencing results to the *E. coli* K12 BW21153 genome.

**FIG S2.**
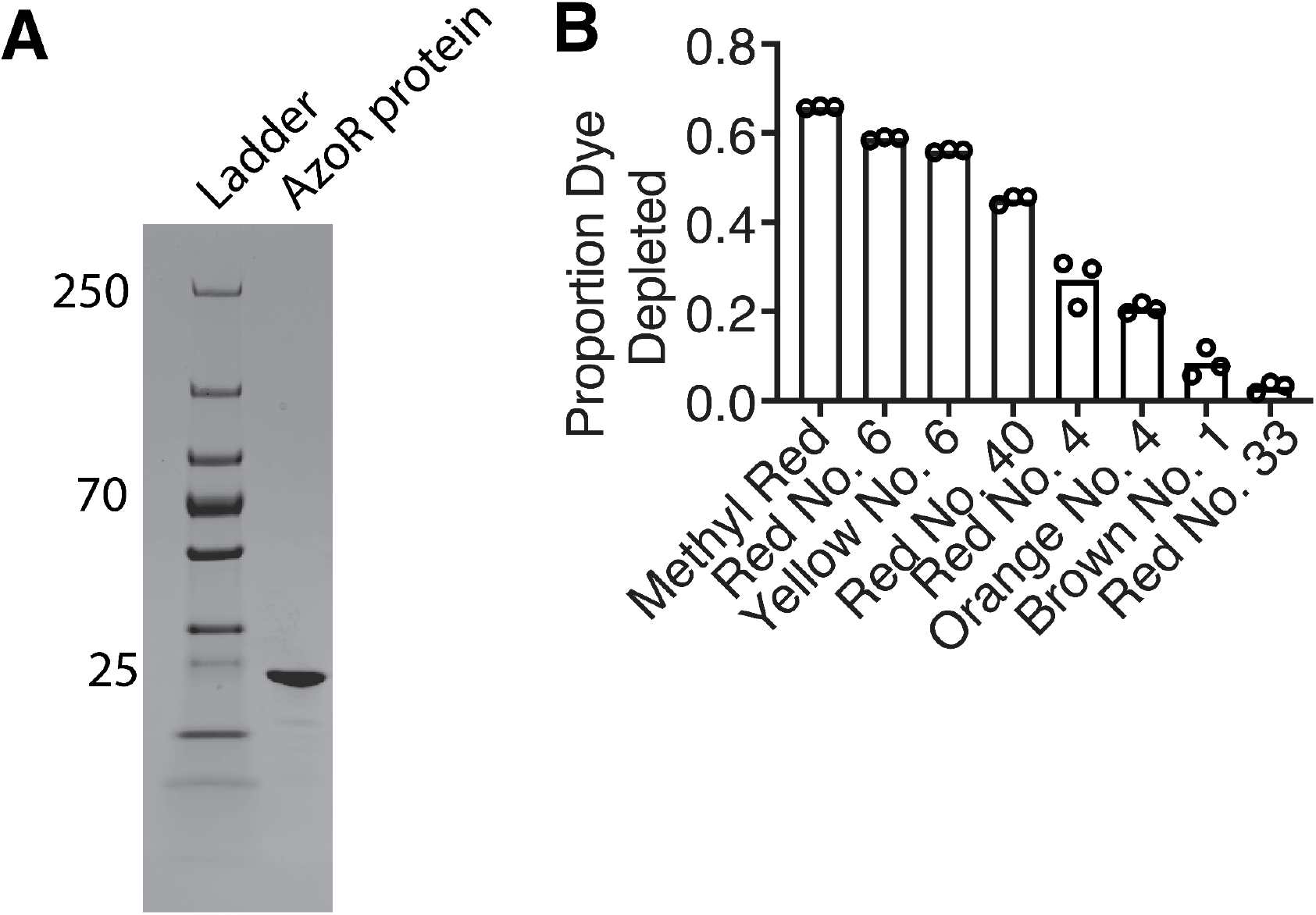
*E. coli* AzoR is sufficient to deplete azo dyes. (A) Purified AzoR protein from *E. coli* AG1 has the expected size (23 kDa) on an SDS-PAGE gel. (B) AzoR protein has variable impacts on the depletion of a panel of FD&C dyes. Incubations were conducted in PBS buffer with 2000μM NADH, and 20μM FMN at 37°C for 15 min, then proportion of the dye depleted was measured by spectrophotometric and calculated relative to no enzyme controls (n=3 biological replicates/strain).

**FIG S3.**
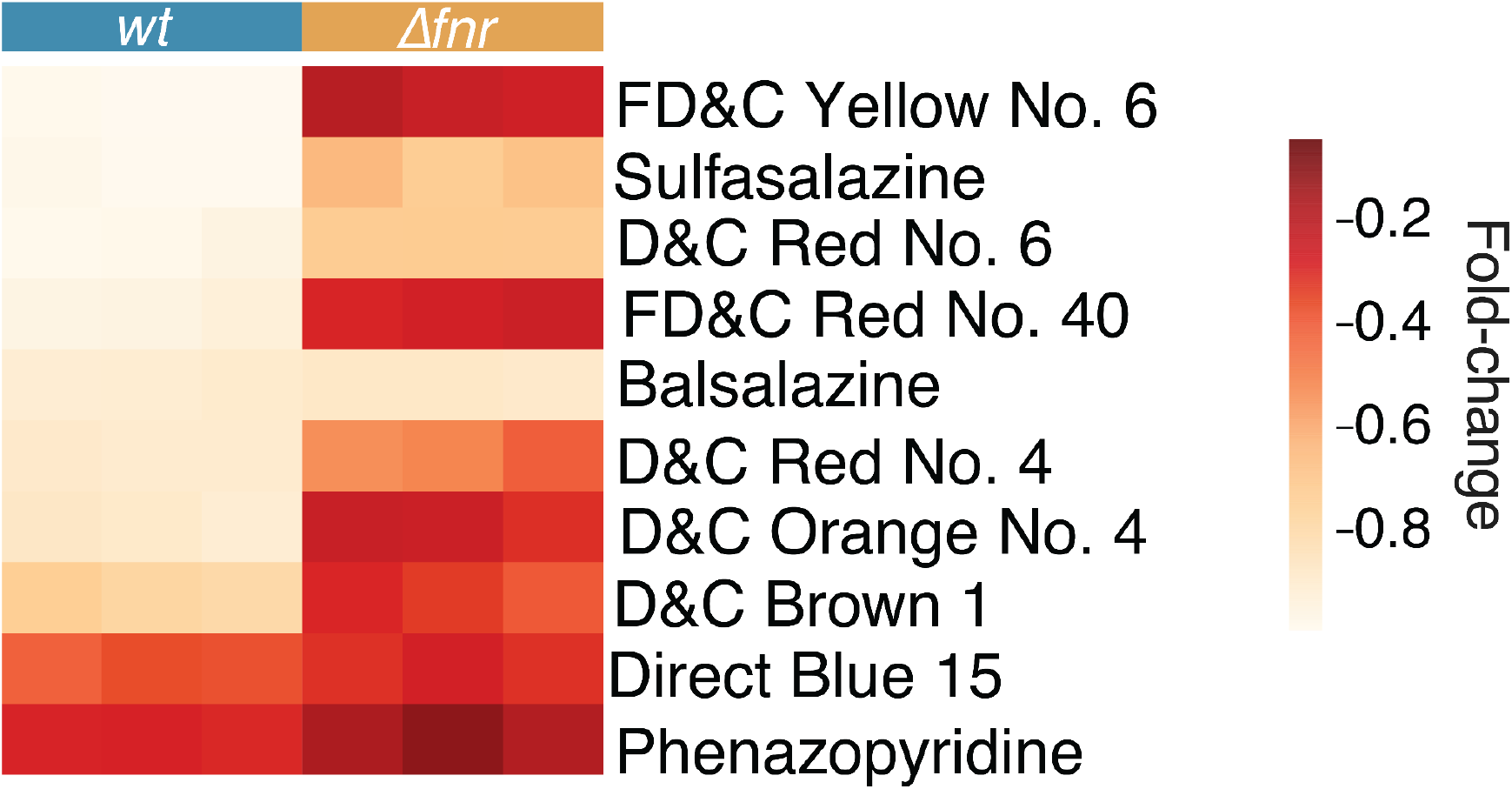
*fnr* is necessary for the depletion of multiple azo dyes. Dye depletion was assessed by spectrophotometry at appropriate wavelengths following 24 hours of incubation in LB media with 4.13 mM L-Cysteine and 250 μM azo dye/drug. Values are fold-change relative to the mean blank concentration (n=3 biological replicates/strain).

**FIG S4.**
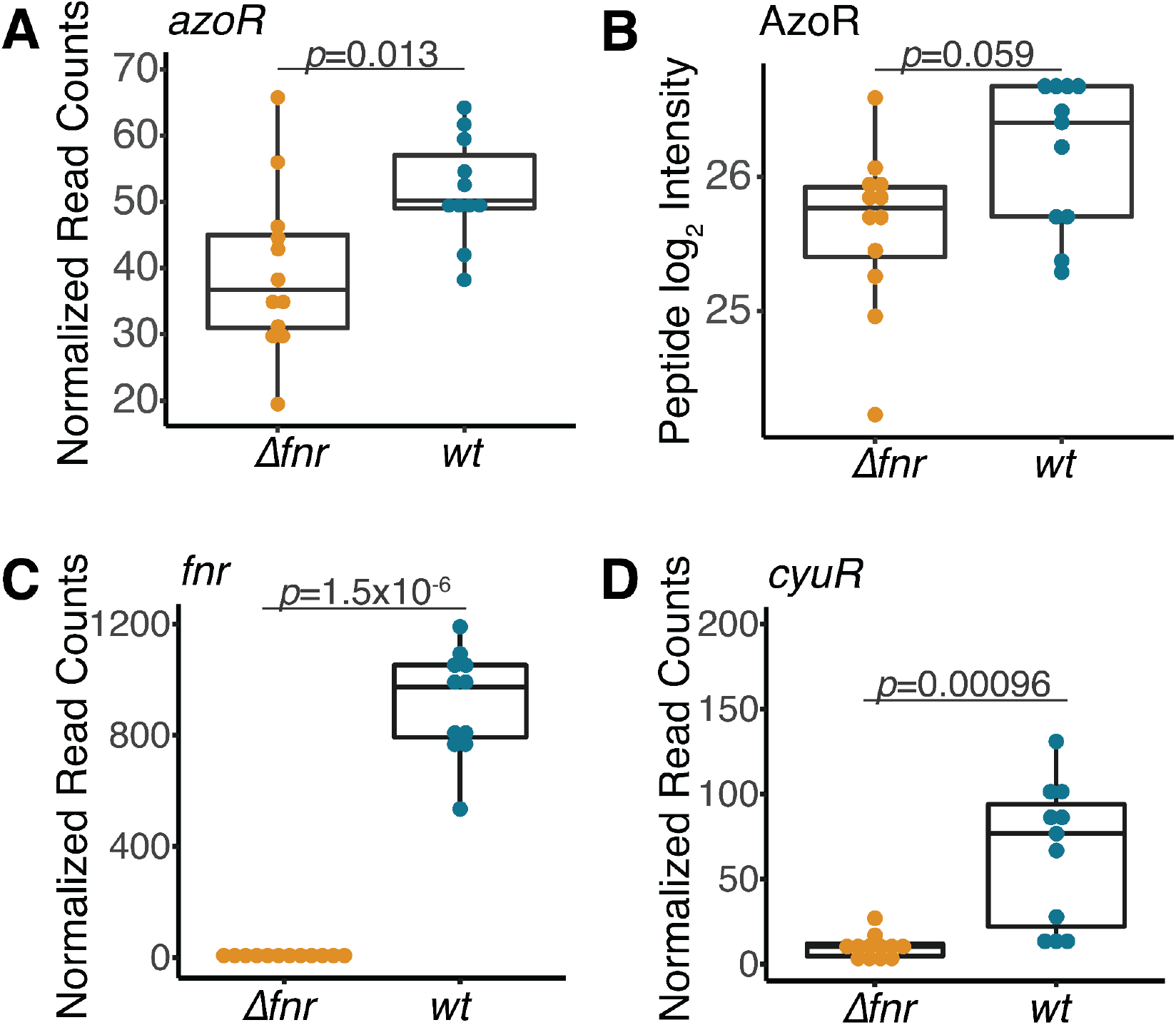
Key transcripts and proteins in *Δfnr E. coli* relative to wild-type controls. (A) *azoR* transcript levels are significantly depleted in the *Δfnr* strain. (B) AzoR peptide levels trend towards lower levels in the *Δfnr* strain. (C,D) *fnr* and *cyuR* transcript levels are significantly lower in the *Δfnr* strain. FNR and cyuR peptides were below the limit of detection. All timepoints and conditions are included within each genotype. *p*-value, Wilcoxon rank sum test. (A,C,D) Normalized read counts are normalized by size factor and have an added pseudo count of 0.5 for log_2_ transformation plotting.

